# Cell-free chromatin particles damage genomic DNA of healthy cells via an ROS-independent mechanism

**DOI:** 10.1101/2024.08.16.608226

**Authors:** Karishma Jain, Gorantla V. Raghuram, Relestina Lopes, Naveen Kumar Khare, Snehal Shabrish, Indraneel Mittra

## Abstract

Several hundred billion cells die daily in the human body, releasing cell-free chromatin particles (cfChPs) in the process, which then enter the blood circulation and get taken up by healthy cells. We have previously reported that, these internalized cfChPs damage genomic DNA as well cause physical damage to mitochondria, resulting in increased mitochondrial ROS production. In the current study, we evaluated the potential damaging effects of the cfChP-induced increase in ROS production on genomic DNA. NIH3T3 mouse fibroblast cells were treated with cfChPs isolated from the sera of healthy individuals (H-cfChPs) or patients with cancer (C-cfChPs) in the presence or absence of the ROS scavenger Mito-TEMPO. The pre-incubation of cfChP-treated cells with Mito-TEMPO abolished ROS production, but did not prevent genomic DNA damage induced by H-cfChPs and C-cfChPs. Our results suggest that cfChP-induced genomic DNA damage occurs in an ROS-independent manner. These findings align with emerging evidence suggesting that mitochondrial ROS may not be a direct cause of genomic DNA damage and suggest that DNA damage attributed to ROS may in-fact be induced by cfChPs. This possibility opens up new therapeutic approaches involving deactivation of cfChPs to retard ageing and other degenerative conditions traditionally attributed to oxidative stress.

## Introduction

Oxidative stress is a major factor regulating ageing and the pathogenesis of various disease conditions^1–6^. Oxidative stress arises from an abundance of reactive oxygen species (ROS) or reactive nitrogen species^7,8^. The levels of ROS, a byproduct of normal cellular metabolism, are kept in check via the antioxidant system. Excessive ROS production disrupts cellular redox equilibrium, leading to mitochondrial dysfunction, protein and lipid oxidation, and DNA damage, further triggering processes such as inflammation and apoptosis^7,9,10^. As maximum cellular oxygen consumption occurs at the mitochondria, these organelles are widely considered as generators of oxidative stress via excess ROS production.

Every day, several hundred billion cells die in the human body^11–12^. In the process, cell-free chromatin particles (cfChPs) are released into the circulation^13^. We have previously demonstrated that circulating cfChPs from dying cells are readily taken up by healthy cells, wherein they integrate into and damage their genomic DNA^14,15^. We have further shown that these cfChPs disrupt mitochondrial structure as well as physically damage their DNA^16^. The latter triggers mitochondrial ROS production, leading to excessive ROS accumulation within the healthy cells^16^.

ROS has been traditionally associated with genomic DNA damage^17–20^. However, experimental evidence demonstrating that mitochondrial ROS release can directly damage genomic DNA is largely lacking^21,22^. As we have previously demonstrated that cfChP treatment of cells leads to abundant ROS production^16^, in this study, we aimed to test whether the cfChP-induced increase in ROS levels contributes to the genomic DNA damage.

## Results

To verify our earlier finding that internalized cfChPs increase mitochondrial ROS production levels within the cell^16^, we treated NIH3T3 mouse fibroblast cells for 4 h with progressively increasing concentrations of cfChPs isolated from the sera of healthy volunteers (H-cfChPs) or patients with cancer (C-cfChPs). The cells were then stained with MitoSOX Red, a dye that detects mitochondrial superoxide radicals (ROS), and analyzed using fluorescent microscopy to estimate mean fluorescence intensity (MFI) per cell. Consistent with our previous observation^16^, H-cfChP and C-cfChP treatments led to a marked increase in ROS levels in the NIH3T3 cells in a dose-dependent manner (Figure 1). As maximum activation of MitoSOX Red was observed with 50 ng of H-cfChPs and 5 ng of C-cfChPs, these concentrations were used in further experiments.

**Figure 1.**
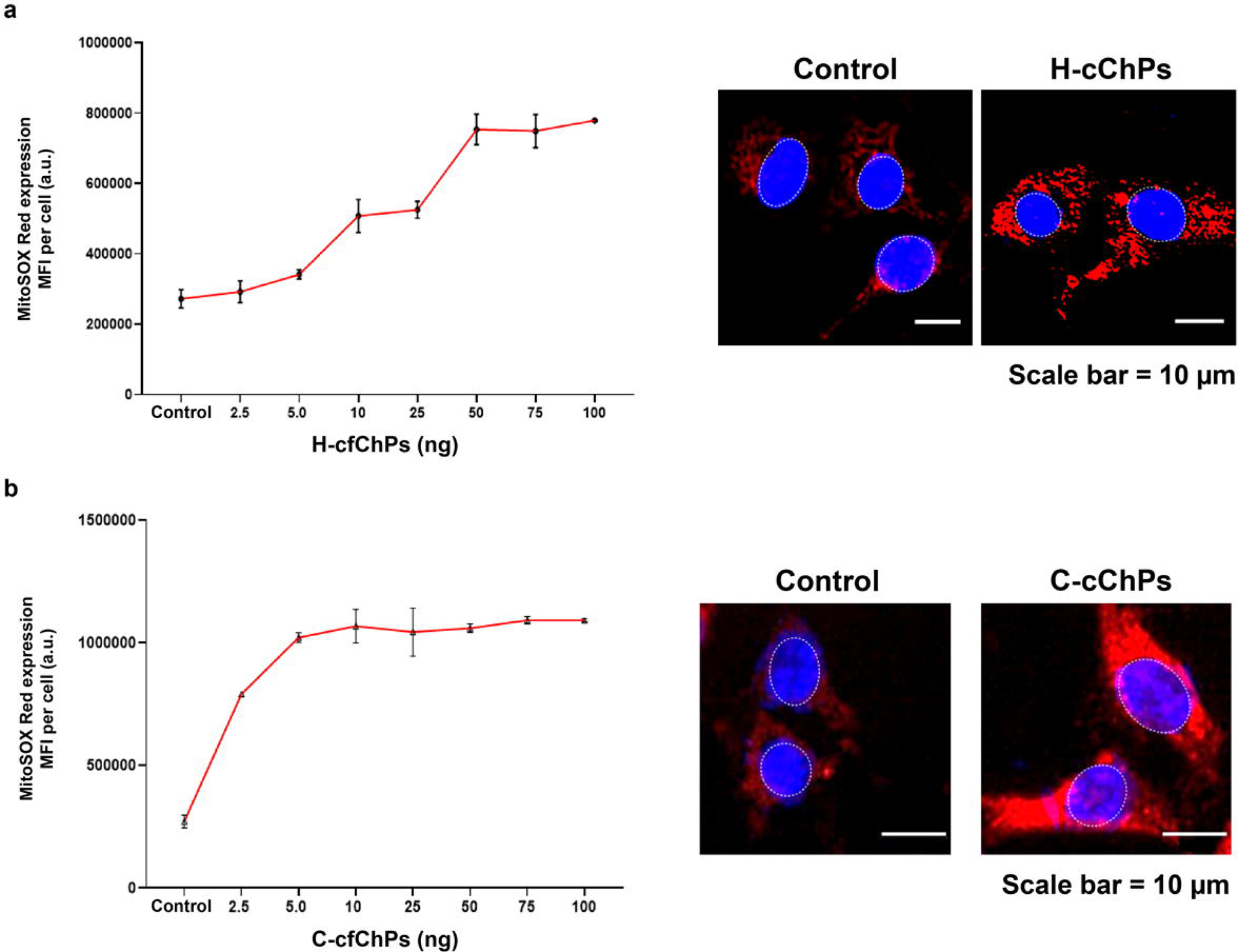
cfChPs activate ROS in a dose-dependent manner. NIH3T3 cells were treated with H-cfChPs and C-cfChPs at progressively increasing concentrations for 4 h. Cells were stained with MitoSOX Red and MFI was estimated from 200 cells after gating and excluding nuclear fluorescence. Dose response curves depicting the activation of MitoSOX red expression in cells treated with (**a**) H-cfChPs and (**b**) C-cfChPs. Results are represented as mean ± SEM values. Representative fluorescence microscopy images of cells treated with H-cfChPs (50 ng; right hand panel in a) and C-cfChPs (5 ng; right hand panel in b) are shown (Scale bar: 10 µm). cfChPs, cell-free chromatin particles; H-cfChPs, cfChPs isolated from the sera of healthy individuals; C-cfChPs, cfChPs isolated from the sera of patients with cancer; MFI, mean fluorescence intensity; a.u., arbitrary units.

Next, we used Mito-TEMPO (MTp), a mitochondrial superoxide radical scavenger, to elucidate the effects of the increased ROS levels in response to the cfChP treatment. NIH3T3 cells were pre-incubated with progressively increasing concentrations of MTp for 1 h, followed by treatment with H-cfChPs (50 ng) or C-cfChPs (5 ng) for 4 h. MTp pre-treatment suppressed the cfChP-induced increase in ROS levels in a dose-dependent manner (Figure 2). The optimum scavenging of MitoSOX Red was observed at concentrations of 50 μM and 75 μM of MTp in H-cfChP- and C-cfChP-treated cells, respectively.

**Figure 2.**
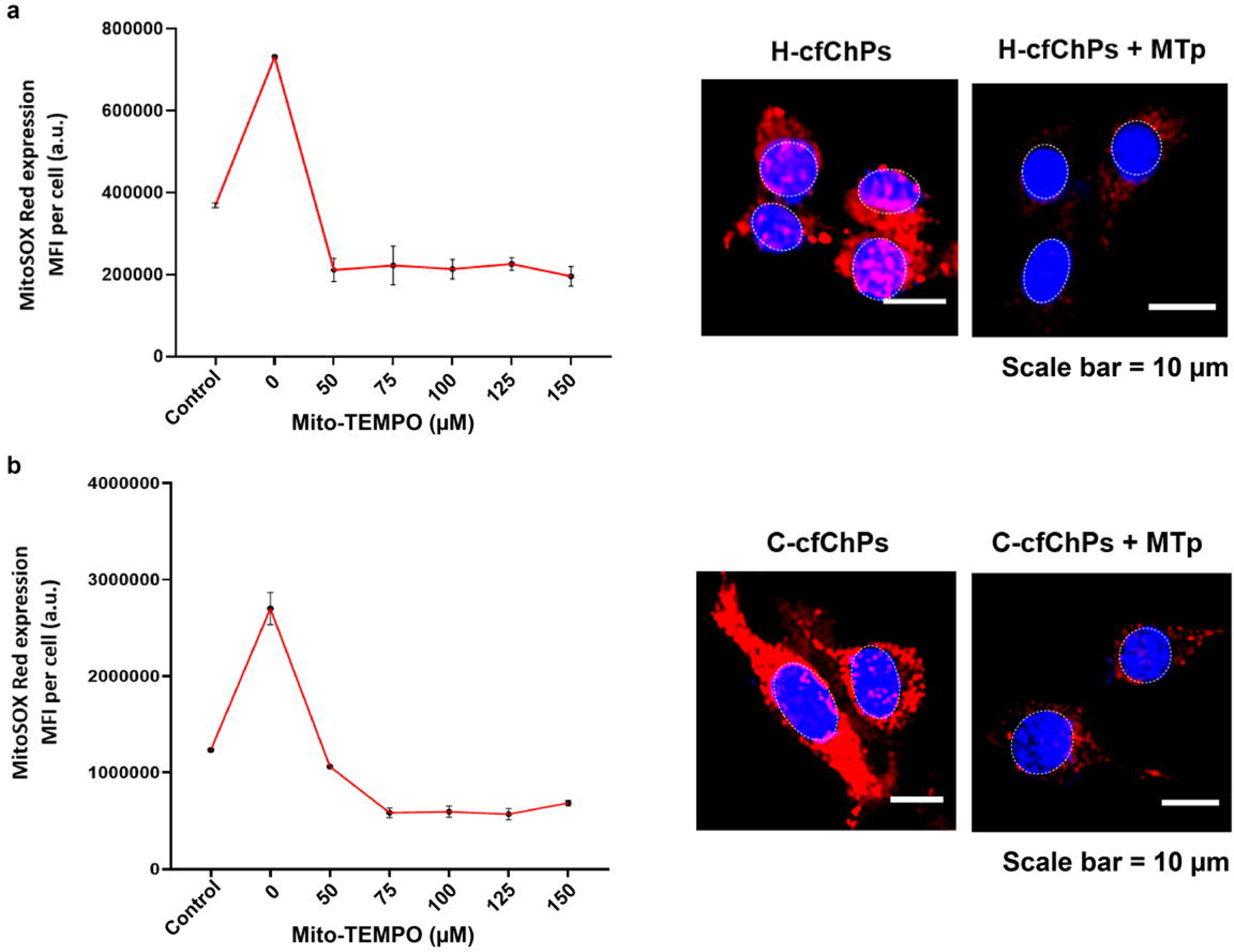
Mito-TEMPO scavenges cfChP-induced ROS in a dose-dependent manner. NIH3T3 cells were pre-incubated with increasing concentrations of Mito-TEMPO for 1 h followed by treatment with (**a**) H-cfChPs (50 ng) and (**b**) C-cfChPs (5 ng) for 4 h. Cells were stained with MitoSOX Red dye and MFI was quantified from 200 cells after gating and excluding nuclear fluorescence. Results are represented as mean ± SEM values. Representative fluorescence microscopy images of cells treated with 50 ng of H-cfChPs and 50 µM of Mito-TEMPO (right hand panel in a) and with 5 ng of C-cfChPs and 75 µM of Mito-TEMPO (right hand panel in b) are shown (Scale bar: 10 µm). cfChPs, cell-free chromatin particles; H-cfChPs, cfChPs isolated from the sera of healthy individuals; C-cfChPs, cfChPs isolated from the sera of patients with cancer; MFI, mean fluorescence intensity; a.u., arbitrary units.

Further, we treated NIH3T3 cells with H-cfChPs (50 ng) or C-cfChPs (5 ng) following pre-incubation with MTp (50 µM and 75 µM MTp for H-cfChP and C-cfChP treatment, respectively). As expected, MTp pre-treatment inhibited ROS production by both forms of cfChPs (Figure 3). Notably, MTp treatment alone did not have any effect on the control NIH3T3 cells (Figure 3). The experiment was done twice with reproducible results.

**Figure 3.**
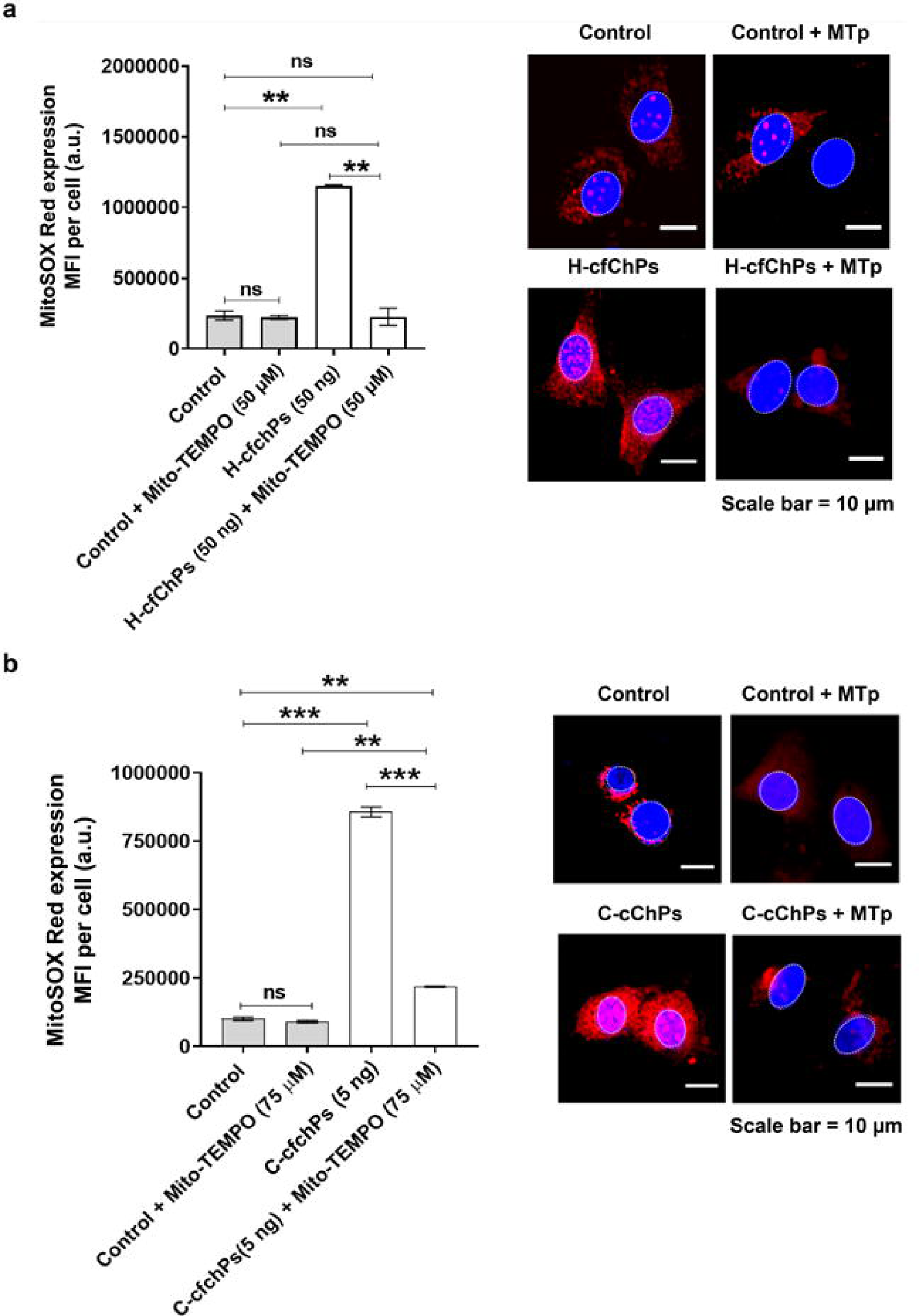
Activation of ROS production in NIH3T3 cells treated with cfChPs in presence and absence of MTp. Histograms illustrating ROS production at 4 h detected by MitoSox Red staining in NIH3T3 cells following treatment with (**a**) H-cfChPs (50 ng) with or without MTp pre-treatment (50 µM), and (**b**) C-cfChPs (5 ng) with or without MTp pre-treatment (75 µM). The MFI was quantified from 200 cells after gating and excluding nuclear fluorescence. Results represent mean ± SEM values. Data were analysed using Student’s *t*-tests. **p<0.01, ***p<0.001, ns = not significant. Representative fluorescence microscopy images of cells treated with 50 ng of H-cfChPs and 50 µM of Mito-TEMPO (right hand panel in a) and with 5 ng of C-cfChPs and 75 µM of Mito-TEMPO (right hand panel in b) are shown (Scale bar: 10 µm). Experiments were performed in duplicate and all images were acquired under the same settings. cfChPs, cell-free chromatin particles; H-cfChPs, cfChPs isolated from the sera of healthy individuals; C-cfChPs, cfChPs isolated from the sera of patients with cancer; MTp, Mito-TEMPO; MFI, mean fluorescence intensity; a.u., arbitrary units. The experiment was done twice with reproducible results.

As we have previously demonstrated that cfChP treatment can induce genomic DNA damage^14,15^, we hypothesized whether the cfChP-induced ROS is responsible for this DNA damage. To test this, we treated NIH3T3 cells with H-cfChPs (50 ng) or C-cfChPs (5 ng) with or without MTp pre-treatment (at 50 µM and 75 µM for H-cfChP and C-cfChP treatment, respectively). Consistent with our previous results, cfChP treatment markedly increased the expression of γH2AX, p-ATM, and p-ATR proteins (Figure 4a–4c), all of which are involved in the DNA damage response (DDR). Notably, cfChP-treated cells with or without MTp pre-treatment showed a similar percentage of cells showing expression of the DDR markers. The experiment was done twice with reproducible results. Our findings suggest that cfChPs induce DDR even in the absence of ROS, indicating that mitochondrial ROS may not account for the genomic DNA damage induced by cfChPs (depicted in an animated illustration; Supplementary Video 1).

**Figure 4.**
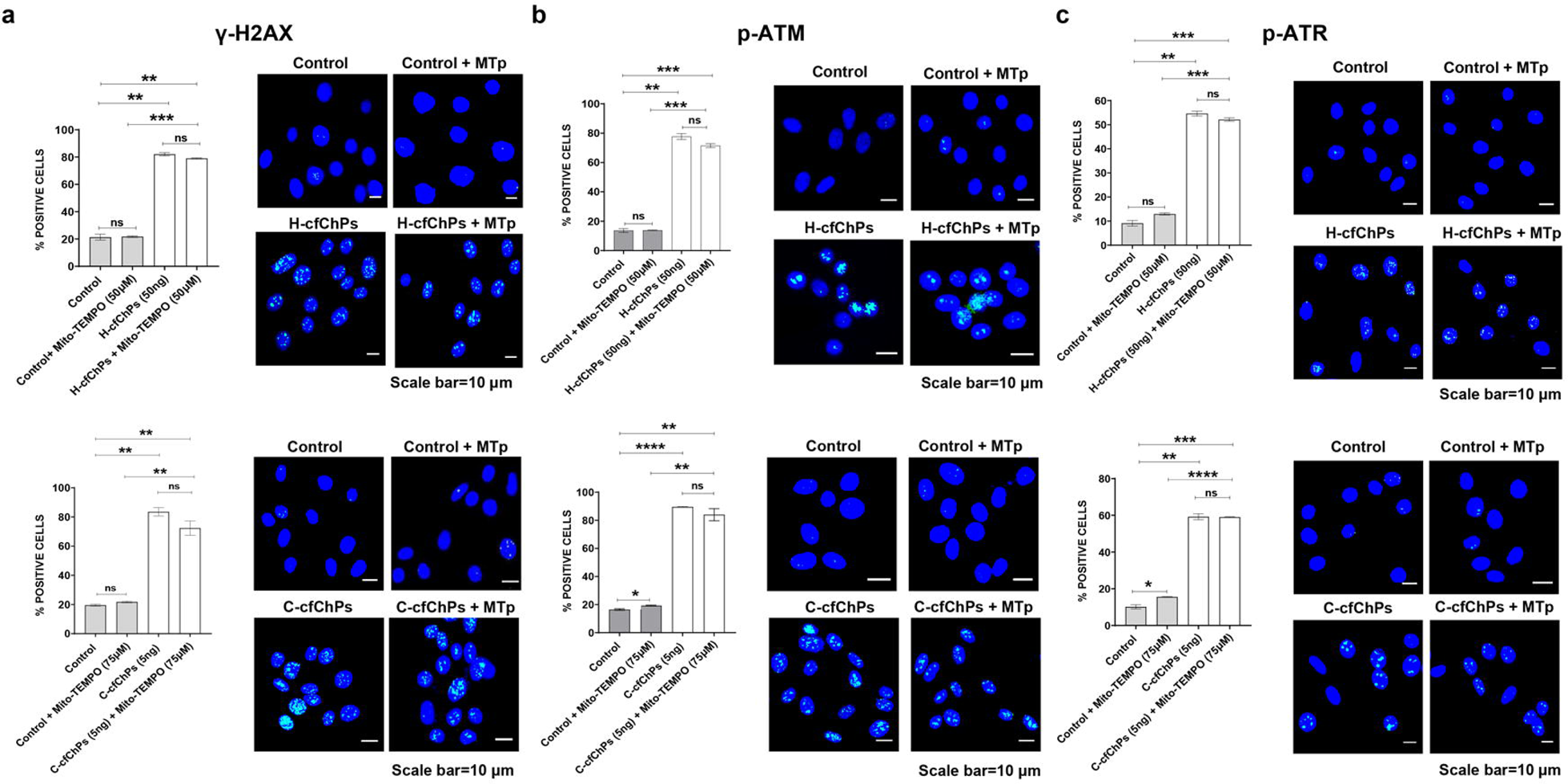
Inhibition of ROS does not prevent genomic DNA damage in cfChP-treated cells. Histograms illustrating the expression of (**a**) γH2AX, (**b**) pATM, and (**c**) pATR after analysing 200 cells and determining the percentage of cells with positive expression signals for the respective proteins. Results are represented as mean ± SEM values. Data were analysed using Student’s *t*-tests. ** p<0.01, *** p<0.001, **** p<0.0001, ns = not significant. Representative images of the expression of (**a**) γH2AX, (**b**) pATM, and (**c**) pATR in untreated control and H-cfChPs (50 ng)- and C-cfChPs (5 ng)-treated cells with and without MTp pre-incubation (50 µM and 75 µM, respectively) are also shown. Scale bars: 10 µm. The histograms and fluorescent images in the upper row are those relating to H-cfChPs and the lower row are those relating to C-cfChPs. Experiments were performed in duplicate and all images were acquired under the same settings. cfChPs, cell-free chromatin particles; H-cfChPs, cfChPs isolated from the sera of healthy individuals; C-cfChPs, cfChPs isolated from the sera of patients with cancer; MTp, Mito-TEMPO. The experiment was done twice with reproducible results.

## Discussion

Our previous studies have demonstrated that circulating cfChPs can enter healthy cells and induce various damaging effects^14–16^, including dsDNA breaks and inflammatory and apoptotic pathways. We have also shown that internalized cfChPs cause a marked increase in ROS levels as well as damage mitochondrial DNA in healthy cells^16^. In the current study, we demonstrate that the cfChP-induced DNA damage does not occur via the increased ROS levels.

H_2_O_2_ treatments have been traditionally used to implicate intracellular ROS in the induction of genomic DNA damage^23–26^. However, the reproducibility of H_2_O_2_ treatments have been extremely difficult, largely because H_2_O_2_ has a short half-life and it is susceptible to enzymatic and chemical elimination in vitro and in cells^27,28^. Furthermore, most studies using H_2_O_2_ to trigger oxidative damage have tended to use concentrations that are disproportionately higher than the physiological concentrations within cells^27^. Consequently, there is a lack of sufficient evidence to demonstrate that the physiological levels of mitochondrial ROS can directly damage nuclear DNA^21,22^. A recent study reported that mitochondrial H_2_O_2_ release does not directly induce genomic DNA damage or mutations and p53 dependent cell cycle arrest, even when used at very high levels compared to what mitochondria normally produce^22^. Ransy et al.^27^ found that experimental exposure to H_2_O_2_ resulted in rapid O_2_ release and drastic reduction in H_2_O_2_ levels within few minutes, thereby explaining why recording of oxidative damage requires the high concentrations found in the literature. Consistent with these reports, our findings suggest that mitochondrial ROS may not significantly contribute to genomic DNA damage.

In the current study, we reconfirmed that cfChPs can induce genomic DNA damage and demonstrated that this damage occurs via an ROS-independent mechanism. Considering that internalized cfChPs accumulate in the nucleus as well as integrate with the genome^14^, the genomic DNA damage may be entirely and directly caused by the interaction of the cfChPs with the genomic DNA. However, further studies are required to confirm this and elucidate the molecular mechanism underlying cfChP-induced genomic DNA damage.

In conclusion, our study shows that cfChPs may be the hitherto unknown agents responsible for the various effects attributed to oxidative stress. Given that several hundred billion to a trillion cells die in the body on a daily basis^11,12^, we hypothesize that circulating cfChPs from dying cells may be the critical physiological triggers that induce mitochondrial DNA damage and ROS production. Therefore, targeting cfChPs and inactivating them could be a promising therapeutic approach to slow down ageing and associated degenerative conditions traditionally linked to oxidative stress. In this context, we have already reported that deactivating cfChPs can retard multiple biomarkers of ageing in C57Bl/6 mice^29^ and downregulate cancer hallmarks and immune check-points in patients with advance oral cancer^30^.

## Methods

### Ethics approval

All study participants provided signed informed consent and the study protocol was approved by the thics Committee of the Advanced Centre for Treatment, Research and Education in Cancer, Tata Memorial Centre (Approval number: 900520).

### Reagents and antibodies

Supplementary Table 1 lists all the names, sources, and catalog numbers of reagents and antibodies used in this study.

### Isolation of cfChPs from the sera of healthy donors and patients with cancer

Blood (10 ml) was collected from five healthy individuals and five patients with cancer; demographic details of the participants are given in Supplementary Table 2. The collected blood samples were allowed to clot before separating the serum for the experiments. To reduce any inter-sample variability, the sera collected from the five healthy individuals and five patients with cancer were pooled for isolating cfChPs. cfChPs were isolated from serum by a method described by us earlier^14^. As reported earlier^14^, electron microscopy revealed the isolated cfChPs to have a beads-on-a-string appearance, which is typical of chromatin. The cfChP concentrations were expressed in terms of their DNA content^14^, which was quantified using the PicoGreen quantification assay (Thermo Fisher Scientific, Waltham, MA, USA), according to the manufacturer’s protocol.

### Cell culture

NIH3T3 mouse fibroblast cells, obtained from the American Type Culture Collection (ATCC, Manassas, VA, USA), were cultured in Dulbecco’s modified Eagle’s medium (DMEM; Gibco, Thermo Fisher Scientific, Catalogue No. 12800-017), supplemented with 10% bovine calf serum (HyClone; Cytiva, Marlborough, MA, USA, Catalogue No. SH30073), at 37°C and 5% CO_2_. Prior to experiments, 6 × 10^4^ cells were seeded onto coverslips in 1.5 ml DMEM and incubated overnight for optimal confluency.

### Estimation of mitochondrial ROS production using MitoSOX Red

#### Dose response analysis of cfChP-induced ROS production

NIH3T3 cells were treated with the isolated H-cfChPs and C-cfChPs at increasing concentrations (viz. 2.5 ng, 5 ng, 10 ng, 25 ng, 50 ng, 75 ng, and 100 ng) and ROS was estimated at 4 h. Briefly, the treated cells were stained with 0.5 µM MitoSOX Red dye in HBSS buffer for 15 min at 37°C, washed with warm buffer, counterstained with Hoechst, and mounted on glass slides. Thereafter, cell-associated fluorescence was measured using a spectral bioimaging system (Applied Spectral Imaging, Israel). The experiments were performed in duplicate and MFI analysis was conducted for 200 cells. As MitoSOX Red generates the by-product mito-ethidine, which intercalates with DNA to generate fluorescence, the nuclei were gated and nuclear fluorescence was excluded from the MFI analysis. The optimum concentration for eliciting ROS production was found to be 50 ng of H-cfChPs and 5 ng of C-cfChPs (Figure 1a and 1b).

#### Dose response analysis of MTp for scavenging ROS production

NIH3T3 cells were pre-incubated with increasing concentrations of MTp (viz. 50 µM, 75 µM, 100 µM, 125 µM, and 150 µM) for 1 h prior to treatment with the optimum H-cfChP (50 ng) and C-cfChP (5 ng) concentrations determined in the above experiment. ROS production was measured 4 h after cfChP treatment by staining with 0.5 µM MitoSOX Red and using the protocol described above. The optimum concentration of MTp required for inhibiting ROS production in response to H-cfChPs and C-cfChPs was found to be 50 µM and 75 µM, respectively (Figure 2a and 2b).

#### Fluorescence microscopy for MFI analysis

Fluorescence microscopy was used to estimate ROS production. Cells were imaged using a spectral bio-imaging system (Applied Spectral Imaging, Israel). For estimation of ROS production based on MFI, images were captured using a 40× air objective. All images were acquired at identical exposure settings to achieve uniform analysis of cell associated fluorescence for both control and treated cells. The MFI was measured using ImageJ (Rasband, W.S., U.S. National Institutes of Health, Bethesda, MD, USA).

### Immunostaining for DDR markers

NIH3T3 cells were pre-incubated for 1 h with 50 µM (in case of H-cfChPs) or 75 µM (in case of C-cfChPs) of MTp, followed by treatment with H-cfChPs (50 ng) or C-cfChPs (5 ng), respectively, for 4 h. Cells were processed for immune-staining for the DDR markers γ-H2AX, p-ATM, and pATR using specific antibodies (listed in Supplementary Table 1). The cells were mounted onto glass slides following counterstaining with Vecta-shield DAPI. Images were acquired using Spectral Bio-Imaging System (Applied Spectral Imaging, Israel). Two-hundred cells were analysed for each DDR marker and the percentage of positively stained cells was calculated.

### Statistical analysis

All data are expressed as mean ± standard error of the mean. Statistical analyses were performed using GraphPad Prism version 8.0.0 (GraphPad Software, San Diego, CA, USA). Data were analysed using Student’s *t*-tests (two-tailed, unpaired) and one-way analysis of variance (ANOVA). Values with p < 0.05 were considered to be statistically significant.

## Acknowledgements

Editorial support, in the form of medical writing, assembling tables and creating high-resolution images based on authors’ detailed directions, collating author comments, copyediting, fact checking, and referencing, was provided by Editage, Cactus Communications. This study was supported by the Department of Atomic Energy, Government of India, through grant CTCTMC to the Tata Memorial Centre awarded to IM.

## Author Contributions

Conceptualization: I.M.; Methodology: K.J., R.V.G., R.L., and N.K.K.; Investigation: K.J. and R.V.G.; Visualization: K.J., R.V.G., and I.M.; Funding acquisition: I.M.; Project administration: R.V.G. and I.M.; Supervision: I.M.; Writing – original draft: R.V.G., S.S., and I.M.; Writing – review & editing: S.S. and I.M.

## References

1. Cui, H., Kong, Y. & Zhang, H. Oxidative stress, mitochondrial dysfunction, and aging. J. Signal Transduct. 2012, 646354 (2012).

2. Itoh, K., Nakamura, K., Iijima, M. & Sesaki, H. Mitochondrial dynamics in neurodegeneration. Trends. Cell Biol. 23, 64–71 (2013).

3. Parish, R. & Petersen, K. F. Mitochondrial dysfunction and type 2 diabetes. Curr.Diab. Rep. 5, 177–183 (2005).

4. Ballinger, S. W. Mitochondrial dysfunction in cardiovascular disease. Free Radic. Biol. Med. 38, 1278–1295 (2005).

5. Boland, M. L., Chourasia, A. H. & Macleod, K. F. Mitochondrial dysfunction in cancer. Front. Oncol. 3, 292 (2013).

6. Yang, J., Luo, J., Tian, X., Zhao, Y., Li, Y. & Wu, X. Progress in understanding oxidative stress, aging, and aging-related diseases. Antioxidants (Basel) 13, 394 (2024).

7. Guo, C., Sun, L., Chen, X. & Zhang, D. Oxidative stress, mitochondrial damage and neurodegenerative diseases. Neural Regen. Res. 8, 2003–2014 (2013).

8. Brand, M. D. The sites and topology of mitochondrial superoxide production. Exp. Gerontol. 45, 466–472 (2010).

9. Nissanka, N. & Moraes, C. T. Mitochondrial DNA damage and reactive oxygen species in neurodegenerative disease. FEBS Lett. 592, 728–742 (2018).

10. Halliwell, B. & Gutteridge, J. M. C. Free Radicals in Biology and Medicine (5^th^ edition). (Oxford university press, 2015)

11. Fliedner, T. M., Graessle, D., Paulsen, C. & Reimers, K. Structure and function of bone marrow hemopoiesis: mechanisms of response to ionizing radiation exposure. Cancer Biother. Radiopharm. 17, 405–426 (2002).

12. Sender, R. & Milo, R. The distribution of cellular turnover in the human body. Nat. Med. 27, 45–48 (2021).

13. Holdenrieder, S., Stieber, P., Bodenmüller, H., Fertig, G., Fürst, H., Schmeller, N., Untch, M. & Seidel, D. Nucleosomes in serum as a marker for cell death. Clin. Chem. Lab. Med. 39, 596–605 (2001).

14. Mittra, I., Khare, N. K., Raghuram, G. V., Chaubal, R., Khambatti, F., Gupta, D., Gaikwad, A., Prasannan, P., Singh, A., Iyer, A., Singh, A., Upadhyay, P., Nair, N. K., Mishra, P. K. & Dutt, A. Circulating nucleic acids damage DNA of healthy cells by integrating into their genomes. J. Biosci. 40, 91–111 (2015).

15. Mittra, I., Samant, U., Sharma, S., Raghuram, G. V., Saha, T., Tidke, P., Pancholi, N., Gupta, D., Prasannan, P., Gaikwad, A., Gardi, N., Chaubal, R., Upadhyay, P., Pal, K., Rane, B., Shaikh, A., Salunkhe, S., Dutt, S., Mishra, P. K., Khare, N. K., Nair, N. K. & Dutt, A. Cell-free chromatin from dying cancer cells integrate into genomes of bystander healthy cells to induce DNA damage and inflammation. Cell Death Discov. 3, 17015 (2017).

16. Raghuram, G. V., Tripathy, B. K., Avadhani, K., Shabrish, S., Khare, N. K., Lopes, R, Pal, K. & Mittra, I. Cell-free chromatin particles released from dying cells inflict mitochondrial damage and ROS production in living cells. Cell Death Discov. 10, 30 (2024).

17. Marnett, L. J. Oxyradicals and DNA damage. Carcinogenesis 21, 361–370 (2000).

18. Dizdaroglu, M., Jaruga, P., Birincioglu, M. & Rodriguez, H. Free radical-induced damage to DNA: mechanisms and measurement. Free Radic. Biol. Med. 32, 1102–1115 (2002).

19. Storr, S. J., Woolston, C. M., Zhang, Y. & Martin, S. G. Redox environment, free radical, and oxidative DNA damage. Antioxid. Redox Signal 18, 2399–2408 (2013).

20. Cadet, J. & Davies, K. J. Oxidative DNA damage & repair: an introduction. Free Radic. Biol. Med. 107, 2–12 (2017).

21. Tripathy, B. K., Pal, K., Shabrish, S. & Mittra, I. A new perspective on the origin of DNA double-strand breaks and its implications for ageing. Genes 12, 163 (2021).

22. van Soest, D. M., Polderman, P. E., den Toom, W. T., Keijer, J. P., van Roosmalen, M. J., Leyten, T. M., Lehmann, J., Zwakenberg, S., Henau, S. D., van Boxtel, R., Burgering, B. M. T. & Dansen, T. B. Mitochondrial H_2_O_2_ release does not directly cause damage to chromosomal DNA. Nat. Commun. 15, 2725 (2024).

23. Hernansanz-Agustín, P. & Enríquez, J. A. Generation of reactive oxygen species by mitochondria. Antioxidants (Basel) 10, 415 (2021).

24. Aruoma, O. I., Halliwell, B., Gajewski, E. & Dizdaroglu, M. Damage to the bases in DNA induced by hydrogen peroxide and ferric ion chelates. J. Biol. Chem. 264, 20509–20512 (1989).

25. Srinivas, U. S., Tan, B. W., Vellayappan, B. A. & Jeyasekharan A. D. ROS and the DNA damage response in cancer. Redox Biol. 25, 101084 (2019).

26. Bertram, C. & Hass, R. Cellular responses to reactive oxygen species-induced DNA damage and aging. Biol. Chem. 389, 211–220 (2008).

27. Ransy, C., Vaz, C., Lombès, A. & Bouillaud, F. Use of H_2_O_2_ to cause oxidative stress, the catalase issue. Int. J. Mol. Sci. 21, 9149 (2020).

28. Marinho, H.S., Cyrne, L., Cadenas, E., & Antunes, F. H_2_O_2_ delivery to cells: Steady-state versus bolus addition. Methods Enzymol 526, 159–173 (2013).

29. Pal, K., Raghuram, G. V., Dsouza, J., Shinde, S., Jadhav, V., Shaikh, A., … & Mittra, I. A pro-oxidant combination of resveratrol and copper down-regulates multiple biological hallmarks of ageing and neurodegeneration in mice. Scientific Reports 12, 17209 (2022).

30. Pilankar, A., Singhavi, H., Raghuram, G. V., Siddiqui, S., Khare, N. K., Jadhav, V., … & Mittra, I. A pro-oxidant combination of resveratrol and copper down-regulates hallmarks of cancer and immune checkpoints in patients with advanced oral cancer: Results of an exploratory study (RESCU 004). Frontiers in Oncology, 12 1000957 (2022).

