## Supplementary Information for "Cell-free chromatin particles damage genomic DNA of healthy cells via an ROS-independent mechanism"

**Supplementary Video 1. An animated illustration showing that cell-free chromatin particles damage genomic DNA by an ROS-independent mechanism**

**Supplementary Table 1**

**Reagents**

| **Sl. No.** | **Reagent** | **Catalogue Number** | **Company** |
| --- | --- | --- | --- |
| 1 | Mito-SOX Red | M36008 | Thermo Fisher Scientific, USA |
| 2 | Mito-TEMPO | SML0737-5MG | Merck-Sigma-Aldrich |
| 4 | Hoechst 33342 | H21492 | Thermo Fisher Scientific, USA |
| 5 | Vecta-Shield DAPI | 101098-042 | Vector Laboratories, USA |
| 7 | Quanti-iT^TM^ PicoGreen®  dsDNA Assay Kit | P7589 | Invitrogen Thermo Fisher Scientific, USA. |

**Antibodies**

| **Sl. No.** | **Antibody** | **Catalogue Number** | **Company** |
| --- | --- | --- | --- |
| 1. | Mouse monoclonal anti- γ-H2AX | 05-636 | Merck, GmbH, Germany |
| 2. | Mouse monoclonal anti-phospho-ATM antibody | 05-740 | Merck, GmbH, Germany |
| 3. | Rabbit anti-ATR polyclonal antibody | ab2905 | Abcam, United Kingdom |
| 4. | Goat Anti-Rabbit IgG H&L (FITC) | ab6717 | Abcam, United Kingdom |
| 5. | Rabbit Anti-mouse (FITC) | AP160F | Merck, GmbH, Germany |

**Supplementary Table 2**

**Healthy Individuals**

| **Sr. No.** | **Age (years)** | **Sex** |
| --- | --- | --- |
| 1. | 36 | Female |
| 2. | 40 | Female |
| 3. | 34 | Male |
| 4. | 42 | Male |
| 5. | 38 | Male |

**Patients with Cancer**

| **Sr. No.** | **Age (years)** | **Sex** | **Diagnosis** | **Stage** |
| --- | --- | --- | --- | --- |
| 1. | 53 | Female | Ca Breast | III |
| 2. | 43 | Female | Ca Cervix | III |
| 3. | 30 | Male | Ca Tongue | III |
| 4. | 76 | Female | Ca Multiple Myeloma | III |
| 5. | 48 | Male | Ca Tongue | IV |
